# From species-area relationships to biodiversity risk assessment

**DOI:** 10.64898/2026.05.15.725338

**Authors:** Marco Tulio Angulo, Serguei Saavedra

## Abstract

Biodiversity is commonly summarized by macroecological mean patterns, most prominently the species-area relationship (SAR) linking habitat area to expected species richness. Yet conservation, policy, and economic decisions increasingly require risk metrics: probabilities of rare but consequential biodiversity shortfalls, including local collapse. Such tail risks are central in finance and insurance but remain difficult to quantify in ecology because the data needed to estimate full richness distributions are rarely available at decision scales. Here we provide a mechanistic route from species-area relationships to biodiversity risk metrics. We show that when regional species abundances are well approximated by Fisher’s log-series, a minimal immigration-extinction mechanism yields a closed-form stationary distribution for local richness whose structure tightly couples the mean SAR to richness variability and lower-tail probabilities. This coupling implies exact fluctuation-response identities and an explicit integral transform that reconstructs collapse probabilities and other tail risk measures directly from the mean SAR. These results define ecological analogues of financial risk metrics—such as collapse probability and lower-tail quantiles—without requiring direct estimation of the full richness distribution. Using high-resolution ForestGEO tree censuses spanning tropical, subtropical, and temperate forests, we find empirical support for these predictions across spatial scales. Together, our results show how widely measurable species-area relationships can be elevated from descriptive averages to operational tools for biodiversity risk assessment and reliability-based conservation planning.

## Introduction

Biodiversity loss has emerged as a systemic risk with ecological, economic, and societal consequences comparable in scale and scope to climate change and financial instability [1–3]. From this perspective, a central question for macroecology and conservation is not only how much biodiversity can an ecosystem sustain on average, but how reliably it avoids dangerous low-diversity states. Reliability is inherently probabilistic: it concerns the likelihood of rare but consequential shortfalls, including local collapse, rather than expected outcomes alone. Macroecology has traditionally addressed biodiversity patterns through expectations and averages [4], most prominently the species-area relationship (SAR) linking habitat area to mean species richness [5–7]. SARs are widely used to guide reserve design and to forecast biodiversity loss under habitat change. Yet a mean is not a risk metric of reliability. Even under broadly similar environmental conditions, births, deaths, and species turnover generate intrinsic stochasticity in local richness [8]. As a result, an area that sustains a target richness on average can still exhibit a substantial probability of falling below that target.

This gap between averages and reliability can be demonstrated using the Barro Colorado Island (BCI) tree census, a 50 ha forest plot in Panama [9]. Suppose a manager seeks to sustain a minimum richness of *Y*_target_ = 4 species within a forest patch. The SAR suggests that an area of *X* = 10.5 m^2^ is sufficient, since the expected richness is *m*(*X*) = 4.07. Yet across patches of this same area, observed richness *Y* ranges from 0 to 12 species (Fig. 1a), yielding a broad probability distribution *π*(*Y* | *X*) of richness outcomes (Fig. 1b). The relevant quantity for decision-making is therefore not the mean *m*(*X*), but the risk of failure Pr(*Y < Y*_target_ | *X*). In the BCI data, this risk is 0.42 at *X* = 10.5 m^2^, implying that more than 40% of patches fall below the target despite meeting it on average. Achieving 90% reliability instead requires roughly doubling area, highlighting how mean-based criteria can substantially underestimate risk.

**Figure 1:**
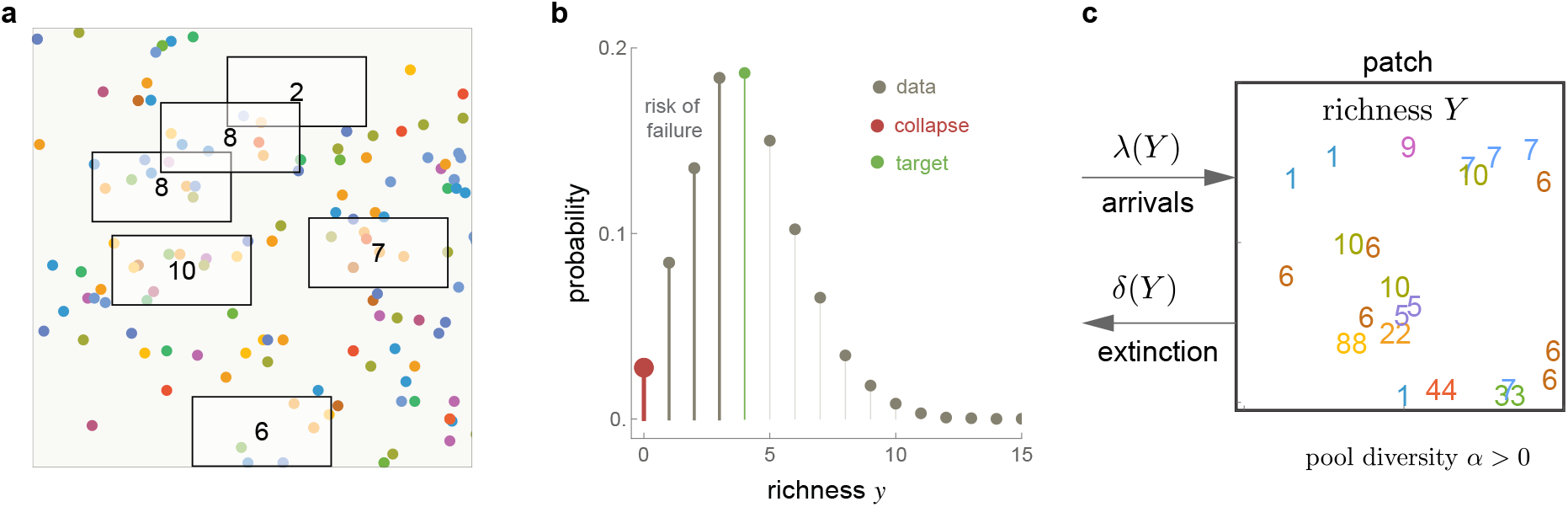
Biodiversity variability and its risks. Figures use experimental census data of trees in a 50 hectare plot in Barro Colorado Island (BCI), Panama [9]. **a**. Example of a 90 m^2^ region of the plot showing individual trees (points) colored by species identity. Rectangles illustrate patches of equal area *X* = 20 m^2^. Numbers inside each rectangle correspond to the observed species richness *Y* in that patch, evidencing its variability. **b**. Empirical distribution *π*(*Y* | *X*) reconstructed from 20,000 random patches of area *X* = 20. This distribution allows calculating biodiversity tail probabilities and associated risk metrics. The first such metric is the probability of biodiversity collapse *π*_0_(*X*) = *π*[*Y* = 0 | *X*] that serves as the lower-tail anchor of the richness distribution (red point). In large forest plots, literal zero-richness states are very unlikely and *π*_0_(*X*) should therefore be interpreted as the theoretical limit of the distributions left tail rather than as an observed state. More broadly, the lower-tail probability P(*Y* ≤*Y*_target_ | *X*) for some small *Y*_target_ could be similarly evaluated. **c**. The inset shows an hypothetical patch with *Y* = 10 species. Numbers inside the patch represent the position of individuals of different species (e.g., there are three individuals of species 1, but only one individual of species 9).

In finance and insurance, such tail risks form the basis of decision-making [10, 11]. Quantities such as ruin probabilities, Value-at-Risk, and Expected Shortfall explicitly characterize the likelihood and severity of rare adverse events [12]. By contrast, ecology lacks comparably tractable tools for quantifying biodiversity tail risk. The challenge is empirical as well as conceptual: estimating the full richness distribution *π*(*Y* | *X*), particularly its lower tail, typically requires dense spatial replication or repeated observations under comparable conditions [13]. Such data are available only for a small number of intensively studied systems such as BCI [14]. In most applications, the mean SAR can be estimated, but the underlying distribution—and therefore risk metrics—remain unidentified.

This creates a fundamental bottleneck. In general, the mean SAR does not determine the richness distribution *π*(*Y* | *X*), and thus cannot uniquely determine the risk of collapse or other low-richness events. A wide family of outcome distributions can be consistent with the same mean curve, implying a wide range of plausible risks. The key question, then, is whether there exist ecological conditions under which this ambiguity collapses: when can a widely measurable mean pattern be promoted to a generator of biodiversity risk metrics?

Here we show that such conditions arise under a classical and empirically common form of regional community structure. When the regional species pool is well approximated by Fisher’s log-series [15], a minimal immigration-extinction mechanism yields a closed-form stationary distribution for local species richness whose structure tightly couples the mean SAR to richness variability and lower-tail probabilities. Fisher’s log-series is a cornerstone of biodiversity theory and is observed across a wide range of ecological systems [16, 17]. Under this structure, the SAR encodes sufficient information to reconstruct tail probabilities through exact fluctuation-response identities and an explicit integral transform.

This mapping from means to tails enables the reconstruction of biodiversity risk metrics directly from species-area relationships. In particular, probabilities of collapse and lower-tail quantiles —ecological analogues of financial risk measures— can be evaluated without direct estimation of the full richness distribution. We refer to this mean-to-tail encoding as biodiversity holography. Using high-resolution ForestGEO tree census data spanning tropical, subtropical, and temperate forests, we find empirical support for these predictions across spatial scales. Together, our results provide a principled route from species-area relationships to biodiversity risk metrics. By identifying conditions under which SARs encode not only expected richness but also variability and extremes, this framework connects macroecology with probabilistic risk analysis and decision making, offering a tractable foundation for reliability-based conservation planning and biodiversity risk assessment.

## Results

### A Fisher log-series pool yields a closed-form richness distribution

We model local species richness *Y* in a habitat patch of area *X* as a continuous-time arrival-extinction process (Fig. 1c). Richness increases through the arrival of novel species at rate *λ*(*Y*) and decreases through extinctions at rate *δ*(*Y*). Consistent with theoretical and empirical studies of spatial scaling [18, 19], we assume that individuals immigrate into the patch at rate *ρX*^*γ*^, where *ρ >* 0 is the baseline arrival rate into a unit-area patch and *γ* ∈ [1*/*2, 1] captures the geometry of immigration, from boundary-controlled (*γ* = 1*/*2) to bulk deposition (*γ* = 1).

When the regional species pool follows Fisher’s log-series with diversity parameter *α >* 0, the probability that an arriving individual belongs to a species not yet present in the patch declines exponentially with local richness (Methods). Specifically, conditional on current richness *Y*, we find that the probability of a novel species is *e*^−*Y/α*^, so richness increases at rate

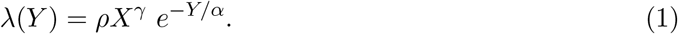

Extinctions occur when the last individual of a species is lost. Let *v >* 0 denote the per-capita mortality rate. Under the same log-series assumption, the expected number of singleton species at richness *Y* is *α*[1 − *e*^−*Y/α*^] (Methods), yielding the extinction rate

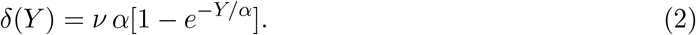

Together, these rates capture ecological saturation: as richness increases, the arrival of novel species slows, while the extinction rate approaches a finite upper bound.

Solving the detailed-balance condition for this birth-death process yields a closed-form for the stationary distribution of richness,

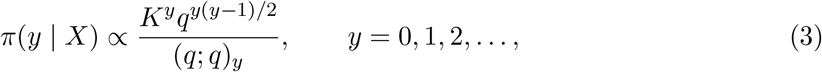

where *q* = *e*^−1*/α*^, 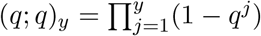 is the *q*-Pochhammer symbol, and

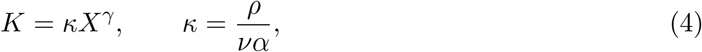

(Methods). We refer to *K* as the *biodiversity potential*, an emergent parameter that summarizes the net balance between immigration, turnover, and area. The immigration turnover ratio *κ >* 0 represents the expected number of immigrants arriving during the average lifetime of an individual. This distribution characterizes the intrinsic stochasticity of richness arising from demographic turnover alone.

Local richness can also vary among patches of equal area due to heterogeneity in local conditions (e.g., permeability, productivity, disturbance). We incorporate this second-order variability by allowing the immigration turnover ratio *κ* to vary lognormally across patches, log *κ*_*i*_ ~ Normal(log *κ, σ*), where the mixing parameter *σ >* 0 characterizes the heterogeneity among patches. The resulting mixed distribution *π*^mix^(*y* | *X*) captures both intrinsic demographic variability and extrinsic spatial heterogeneity, a key contributor to biodiversity risk.

### From richness distributions to biodiversity risk metrics

The stationary distribution in Eq. (3) directly defines biodiversity risk metrics. Of particular interest is the probability of biodiversity collapse,

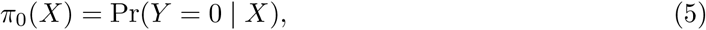

which anchors the lower tail of the richness distribution. More generally, lower-tail probabilities Pr(*Y* ≤ *Y*_target_ | *X*) quantify the likelihood that richness falls below ecologically meaningful thresholds. These quantities are direct analogues of ruin probabilities and lower-tail quantiles used in finance and insurance to assess downside risk [11, 12].

In practice, however, estimating such tail probabilities appears to require direct knowledge of *π*(*Y* | *X*), which in turn demands dense spatial replication or long time series rarely available outside a handful of intensively studied ecosystems. The next section shows that the structure of Eq. (3) resolves this apparent impasse by tightly coupling the mean species-area relationship to the entire richness distribution.

### Structural predictions linking means, variability, and risk

The stationary distribution admits an exact representation as a sum of independent Bernoulli trails,

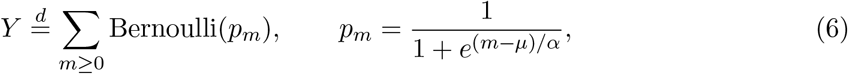

with *µ* = *α* log *K* (Methods and Box 1). This representation reveals that richness behaves as the number of occupied “ecological modes”, each mode *m* filled with a logistic probability *p*_*m*_. Mathematically, this structure is identical to the Fermi–Dirac distribution of statistical physics [20], but here it emerges solely from ecological immigration-extinction dynamics rather than from energetic constraints. Eq. (6) implies a sharp occupancy threshold at *m* = *µ* (Box 1a). Modes below this threshold are almost always filled (i.e., *p*_*m*_ *>* 1*/*2), whereas modes above it are rarely occupied (i.e., *p*_*m*_ *<* 1*/*2). Richness therefore accumulates rapidly up to a level set by regional diversity and immigration pressure *µ* = *α* log *κ* + *αγ* log *X*, and then increases only intermittently as higher modes flicker in and out.

The structure of Eq. (6) yields three key predictions that link the mean SAR to biodiversity risk metrics (Box 1).

i. **Unified scaling with area**. The mean richness follows

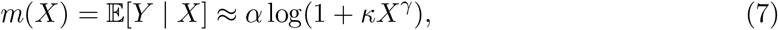

which behaves as a power law at small areas and becomes asymptotically logarithmic at large areas (Fig. 2a,b). In parallel, the variance Var(*Y* | *X*) increases sigmoidally with area and saturates at *α*(1 + *ασ*^2^) (Fig. 2c). These scalings imply diminishing returns of area for both mean richness and risk reduction once ecological saturation is reached.
ii. **Fluctuation–response identity**. Because occupancy probabilities are logistic, changes in mean richness with area determine variability:

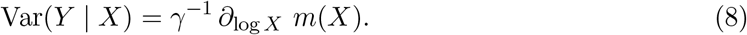 Thus, on a semilogarithmic area axis, the local slope of the SAR is proportional to richness variance (Box 2b). This identity turns the mean SAR into a quantitative proxy for uncertainty and volatility in richness outcomes.
iii. **Mean-to-tail reconstruction (“biodiversity holography”)**. Most importantly for risk assessment, the mean SAR determines the full richness distribution, including its lower tail. In particular, the probability of biodiversity collapse satisfies

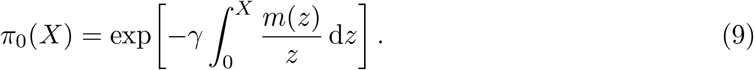

**Figure 2:**
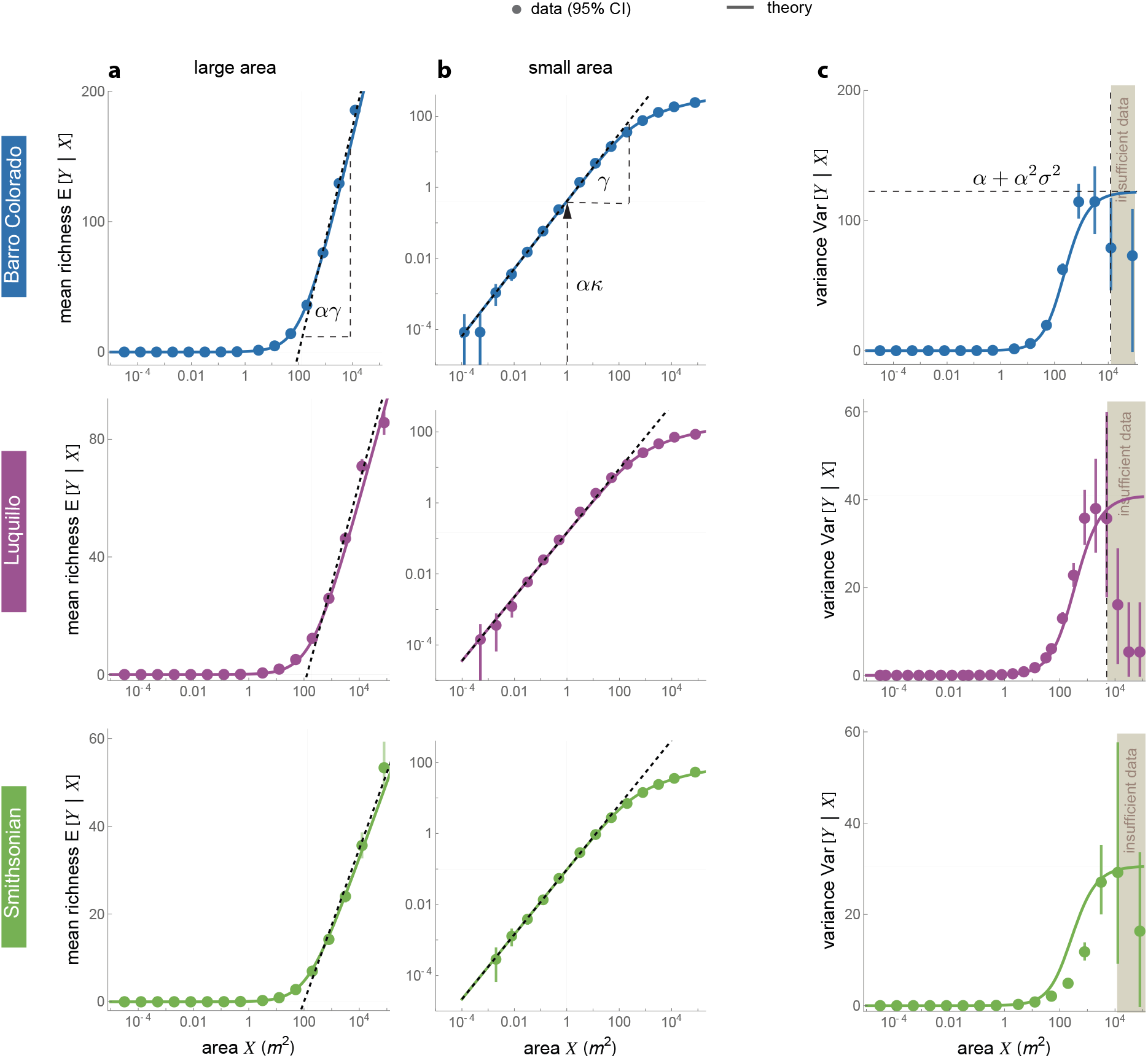
Mean and variance of species richness across spatial scales. Circles with whiskers denote ForestGEO tree census data with 95% confidence intervals. Solid lines show theoretical predictions of the stationary distribution with parameters inferred only from the first two moments (Table 1). Colors correspond to three different plots (Barro Colorado in Panama, Luquillo in Puerto Rico, and Smithsonian in the USA). **a**. For large areas, the model predicts 𝔼[*Y*|*X*] ≈*α* log(*κ*) + *αγ* log *X*. In semi-log coordinates, richness therefore increases linearly with slope *αγ*, which is estimated from the dashed black line. **b**. For small areas, richness follows the classic power-law form 𝔼 [*Y*|*X*] ≈*ακX*^*γ*^. In loglog space, the slope (*γ*) and intercept (*ακ*) determine the effective immigration scaling and regional diversity (dashed black line). **c**. The variance satisfies Var[*Y*|*X*] ≈ *v*(*X*)[1 + *σ*^2^*v*(*X*)], where *v*(*X*) = *ακX*^*γ*^*/*(1 + *κX*^*γ*^). Variance saturates at *α* + *α*^2^*σ*^2^, allowing estimation of the heterogeneity parameter *σ*. At large areas (right of dashed vertical line), estimates become unreliable because few independent patches remain. This is a structural limitation of all spatial datasets.

**Table 1:** Inferred parameters. Smithsonian refers to the Smithsonian Conservation Biology Institute.

| ecosystem | Fisher's diversity $\alpha$ | turnover ratio $\kappa$ | allometric exponent $\gamma$ | mixing $\sigma$ |
| --- | --- | --- | --- | --- |
| Barro Colorado | 39.4767 | 0.0101229 | 0.949571 | 0.230224 |
| Luquillo | 16.4298 | 0.00919816 | 0.908344 | 0.299531 |
| Smithsonian | 8.29032 | 0.011523 | 0.9111779 | 0.569777 |

More general lower-tail probabilities and quantiles follow from an explicit Fourier/Cauchy inversion of the probability generating function (Box 1 and Methods). This mapping allows biodiversity risk metrics to be evaluated directly from the mean SAR, without fitting the full distribution.

As biodiversity potential *K* declines, the theory predicts a sharp increase in collapse risk below a characteristic threshold *K*\*, defined implicitly by Li_2_(−*K*\*) = −*α*^−1^ log 2. Because *K* = *κX*^*γ*^, this corresponds to a critical area *X*\* below which biodiversity collapse is no longer a rare event but a likely outcome.

### Empirical support across forest ecosystems

We tested these predictions using tree census data from three independent ForestGEO plots spanning tropical (Barro Colorado Island, Panama), subtropical (Luquillo, Puerto Rico), and temperate (Smithsonian Conservation Biology Institute, USA) forests. These plots provide spatially explicit censuses of all stems above a diameter threshold, allowing richness to be measured across nested spatial grains and enabling empirical estimation of mean richness, variance, and full distributions as functions of area (Methods and Supplementary Fig. 1).

Across all three forests, mean richness exhibits the predicted transition from power-law scaling at small areas to slower, logarithmic growth at larger areas (Fig. 2a,b). Richness variance increases sigmoidally and approaches the predicted plateau set by regional diversity and spatial heterogeneity (Fig. 2c). Parameter estimates obtained using only the first two moments recover both the observed mean and variance across spatial scales (Table 1).

Using these parameters, the model accurately reproduces the full richness distribution at fixed areas, including the lower tail (Fig. 3a), without additional fitting. We then tested the central risk prediction: whether biodiversity collapse probabilities can be reconstructed from the mean SAR alone. The holographically reconstructed *π*_0_(*X*), computed directly from the empirical mean curve using Eq. (9), closely matches the observed frequency of zero-richness subplots across areas and across all three forests (Fig. 3b). The same approach accurately predicts lower-tail quantiles, including the 5th-percentile richness (Biodiversity-at-Risk), across ecosystems and spatial scales (Fig. 3c).

**Figure 3:**
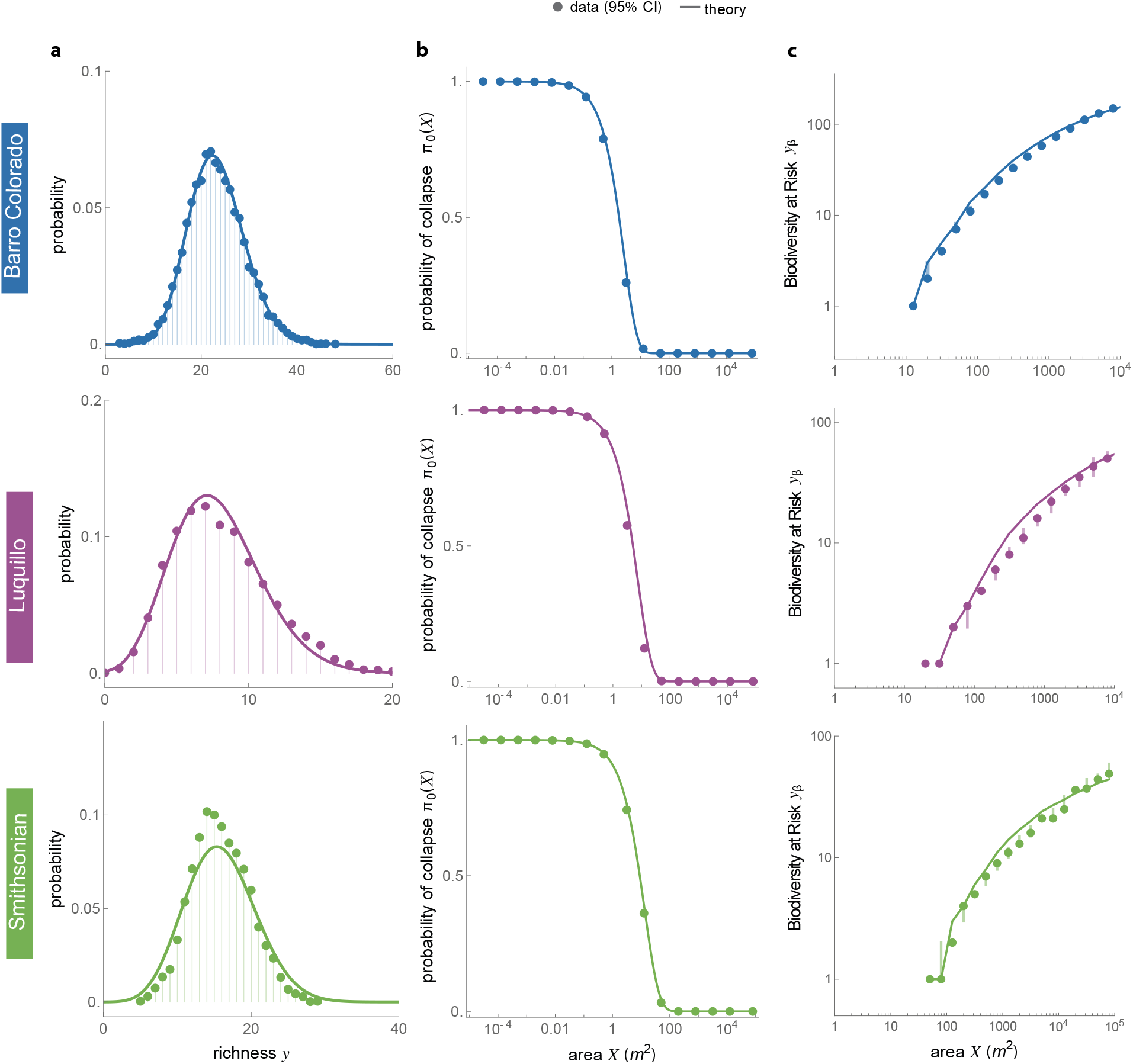
Theoretical predictions match observed biodiversity variability and tail risks. Circles and whiskers show ForestGEO data with 95% confidence intervals. Lines show theoretical predictions using only the parameters inferred from the mean and variance in Fig. 2. No additional fitting is performed for the full distribution or any tail-risk quantity. **a**. Full distribution *π*(*Y* |*X*) of richness at fixed area (*X* = 100 for Barro Colorado and Luquillo, *X* = 1000 for Smithsonian) **b**. Probability of biodiversity collapse *π*_0_(*X*) reconstructed from the mean richness curve𝔼 [*Y* |*X*] ≈*m*(*x*) = *α* log(1+*κX*^*γ*^) using holography. **c**. Biodiversity-at-Risk *y*_0.05_, the worst richness value exceeded only 5% of the time. The predictions accurately recovers the empirical 5th-percentile richness across forests and scales, demonstrating that biodiversity tail risks are encoded in the mean speciesarea curve.

Together, these results support the central claim of the framework: under empirically testable conditions, species-area relationships encode sufficient information to evaluate biodiversity risk metrics, enabling a direct translation from macroecological means to probabilistic assessments of biodiversity collapse and shortfall risk.

## Discussion

Our results establish a direct and mechanistic connection between a classical macroecological pattern —the species-area relationship— and probabilistic measures of biodiversity risk. Under empirically testable conditions, a widely measurable mean curve can be promoted to a generator of risk metrics, including the probability of collapse and lower-tail quantiles of species richness.

This provides a principled route from descriptive averages to reliability-based assessments, addressing a long-standing gap between macroecological theory and decision-relevant measures of biodiversity risk.

A central insight of the framework is that biodiversity risk is governed by the *distribution* of richness outcomes rather than by their expectation alone. Species-area relationships summarize how richness accumulates with area on average, but they do not, in general, determine the likelihood of rare but consequential shortfalls. The results here identify conditions under which this limitation is overcome. When regional species abundances are well approximated by Fisher’s log-series and local dynamics are dominated by immigration-extinction turnover, the stationary richness distribution acquires a constrained statistical structure. This structure tightly links mean richness, variability, and lower-tail probabilities, allowing risk metrics to be reconstructed directly from the SAR without direct estimation of the full distribution. In particular, the predicted scaling for expected richness of Eq. (7) bridges the classical power-law SAR at small scales [21] with the slower logarithmic growth reported in large forest plots [22], providing a mechanistic route to multi-phase SAR shapes [17, 23].

From a risk perspective, this mirrors developments in finance and insurance, where tail probabilities and quantiles —rather than means— form the basis of decision-making. Quantities such as ruin probabilities, Value-at-Risk, and Expected Shortfall are used to characterize exposure to rare adverse events and to set reliability targets [11, 12]. The biodiversity risk metrics derived here are ecological analogues of these concepts. Collapse probabilities and lower-tail quantiles summarize the likelihood and severity of low-richness outcomes across spatial scales, providing interpretable measures of downside risk that are directly relevant to conservation planning, reserve design, and emerging biodiversity risk assessments. This is potentially relevant for emerging nature-related assessment and disclosure efforts that call for quantifiable measures of downside ecological outcomes [3, 24].

Mathematically, the tractability of the framework arises from the fact that the stationary richness distribution factorizes into independent logistic occupancies, a structure identical to the Fermi-Dirac distribution in statistical physics [20]. In the ecological context, this structure reflects saturation under turnover: a small number of occupancy modes are almost always filled, corresponding to persistent taxa, while a long tail of higher modes flickers in and out as rare or transient species. This constrained occupancy spectrum explains why richness variance saturates with area and why collapse risk responds sharply once biodiversity potential falls below a characteristic threshold. Importantly, this physics analogy is not merely metaphorical: it provides exact identities linking the mean SAR to variance and tail probabilities, which underpin the mean-to-tail holographic reconstruction central to the framework.

Our framework is closely related to, but distinct from, classical neutral theory. Neutral models emphasize demographic equivalence and immigration from a regional pool, and they have been highly successful at explaining species abundance distributions, including Fisher’s log-series [25]. However, neutral theory typically focuses on abundance patterns and expected richness, rather than on the stationary distribution of species richness itself. As a result, neutral formulations do not generally yield closed-form expressions for richness variability or lower-tail probabilities, and therefore do not provide direct access to biodiversity risk metrics. By contrast, the present framework derives the full stationary distribution of local richness under a Fisher-consistent immigrationextinction process, making it possible to quantify collapse probabilities and lower-tail quantiles explicitly. In this sense, our results complement neutral theory by extending its demographic logic from abundance patterns to probabilistic assessments of richness reliability.

The empirical results from ForestGEO plots spanning tropical, subtropical, and temperate forests support the theoretical predictions across a broad range of spatial scales. Using only the first two moments of richness, the model reproduces the full richness distribution, including its lower tail, and accurately reconstructs collapse probabilities and lower-tail quantiles directly from the observed mean SAR. These findings suggest that, in these systems, the assumptions underlying the framework —approximate stationarity, dominance of immigration-extinction turnover, and a log-series regional pool— are sufficiently well satisfied for biodiversity risk metrics to be meaningfully inferred from speciesarea data.

The framework also yields qualitative insights into conservation trade-offs. Because collapse probability decreases monotonically with area and its logarithm is a convex function of log-area, habitat fragmentation increases collapse risk within individual patches (Methods). At the same time, unequal fragment sizes can reduce the probability of simultaneous collapse across all patches, revealing a diversificationconcentration trade-off analogous to portfolio effects in finance. Such results illustrate how probabilistic risk metrics can clarify questions that are ambiguous when considered solely in terms of mean richness.

Several limitations delineate the scope of the present results. First, the framework assumes an approximately stationary immigration-extinction balance. Ecosystems undergoing strong successional change, directional land-use shifts, or rapid climatic forcing may violate this assumption, limiting the applicability of stationary risk metrics. Second, species interactions and density-dependent feedbacks enter only implicitly through effective arrival and extinction rates.

Systems dominated by strong priority effects or alternative stable states may exhibit richness distributions that deviate from the form derived here. Third, the reconstruction relies on the regional species pool being close to Fisher’s log-series, particularly in its rare-species tail. Systematic departures from this structure are expected to modify the occupancy spectrum and, consequently, tail-risk estimates. Finally, spatial autocorrelation and dispersal kernels are not explicitly modeled and may inflate variance at intermediate spatial scales.

Despite these limitations, the framework provides clear diagnostics for applicability. Empirically, its use is best justified when (i) regional abundance distributions are approximately log-series, (ii) richness-area curves and their variance are stable across comparable patches, and (iii) spatial autocorrelation does not dominate richness fluctuations at the sampling grain. Where these conditions hold, species-area relationships can be elevated from descriptive summaries to quantitative tools for biodiversity risk assessment.

More broadly, the results suggest a pathway for integrating macroecology with probabilistic risk analysis. By translating species-area relationships into biodiversity risk metrics, the framework enables reliability targets to be formulated and compared across landscapes using data that are already widely collected. This opens the door to biodiversity stress testing analogous to practices in finance, where exposure to rare but consequential events is evaluated under explicit assumptions. Extending this approach to other biodiversity dimensions, such as functional or phylogenetic diversity, and to non-stationary settings represents a promising direction for future work.

In summary, this study shows how species-area relationships can be systematically mapped into biodiversity risk metrics under well-defined ecological conditions. By linking macroecological means to probabilistic extremes, the framework provides a tractable foundation for reliability-based conservation planning and for the broader integration of biodiversity into risk-oriented environmental decision-making.

### BOX 1. The structure of the stationary distribution and its implications.

The stationary distribution of richness factorizes exactly as a sum of independent Bernoulli trials (Methods), implying that

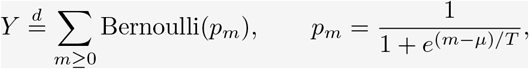

where *µ* := *α* log *K* and *T* := *α*. Notably, this representation is equivalent to the Fermi-Dirac distribution in quantum physics [20], with *µ* representing the chemical potential and *T* the effective temperature (measured directly in energy units where Boltzmann constant equals 1). This structure implies several useful properties (see Methods for proofs):

1. **Simple scaling of the first two moments**. The closed-form moment identities 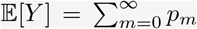 and 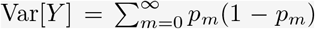. Using a Euler-MacLaurin approximation replacing sums by integrals we obtain:

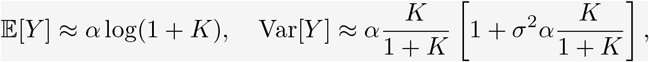

where *K* = *κX*^*γ*^. For large areas *X* → ∞ we obtain 𝔼 [*Y*] ≈ *α* log(*κX*^*γ*^) = *α* log *κ* + *αγ* log *X*. Therefore, in a semi-log plot of 𝔼 [*Y* | *X*] as a function of log *X*, mean richness should increase linearly with slope *αγ*. For small areas *X* → 0 where log(1 + *κX*^*γ*^) ≈ *κX*^*γ*^, the model reduces to the classic allometric speciesarea relation 𝔼 [*Y*] ≈ *ακX*^*γ*^. In a log-log plot of log 𝔼 [*Y*] as a function of log *X*, richness should increase linearly with slope *γ* and intercept *ακ*. The mean richness at *X* = 1 and its slope for large areas and small areas give three observations from which the parameters *α, γ* and *κ* can be inferred. Skewness is positive for small *K* and decays like 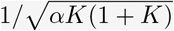with heterogeneity increasing right-skew near *K* ≈ 1.
2. **A generator determines the full distribution**. All cumulants of the richness distribution are determined by the risk “free energy” *F* (*µ, T*) = − log *π*_0_(*µ, T*) as

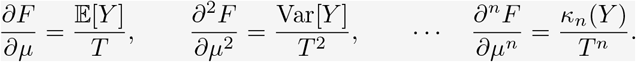 This has several implications. For example, using the first two equation yields Var[*Y*] = *T ∂*_*µ*_E[*Y*] or equivalently Var[*Y*] = *γ*^*−*1^ *∂*_log *X*_ 𝔼[*Y*]. That is, in a semi-log plot of 𝔼[*Y*| *X*] as a function of log *X*, the slope at *X* equals *γ*Var[*Y*] (panel b).
3. **Holography from mean richness**. The mean richness curve *m*(*X*) = 𝔼 [*Y* |*X*] fully characterizes any tail risk ℙ [*Y* ≤ *y*] via the Fourier/Cauchy inversion formula

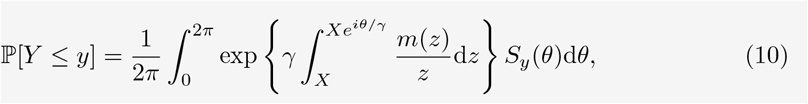

where *S*_*y*_(*θ*) := *e*^*−iyθ/*2^ sin (*y* + 1)*θ/*2 */* sin(*θ/*2). For example, taking *y* = 0, panel c compares the exact and holographic prediction of the probability of biodiversity collapse *π*_0_(*X*) = *π*(*Y* = 0|*X*) using the approximation *m*(*X*) ≈*α* log(1 + *κX*^*γ*^), showing they are practically indistinguishable.

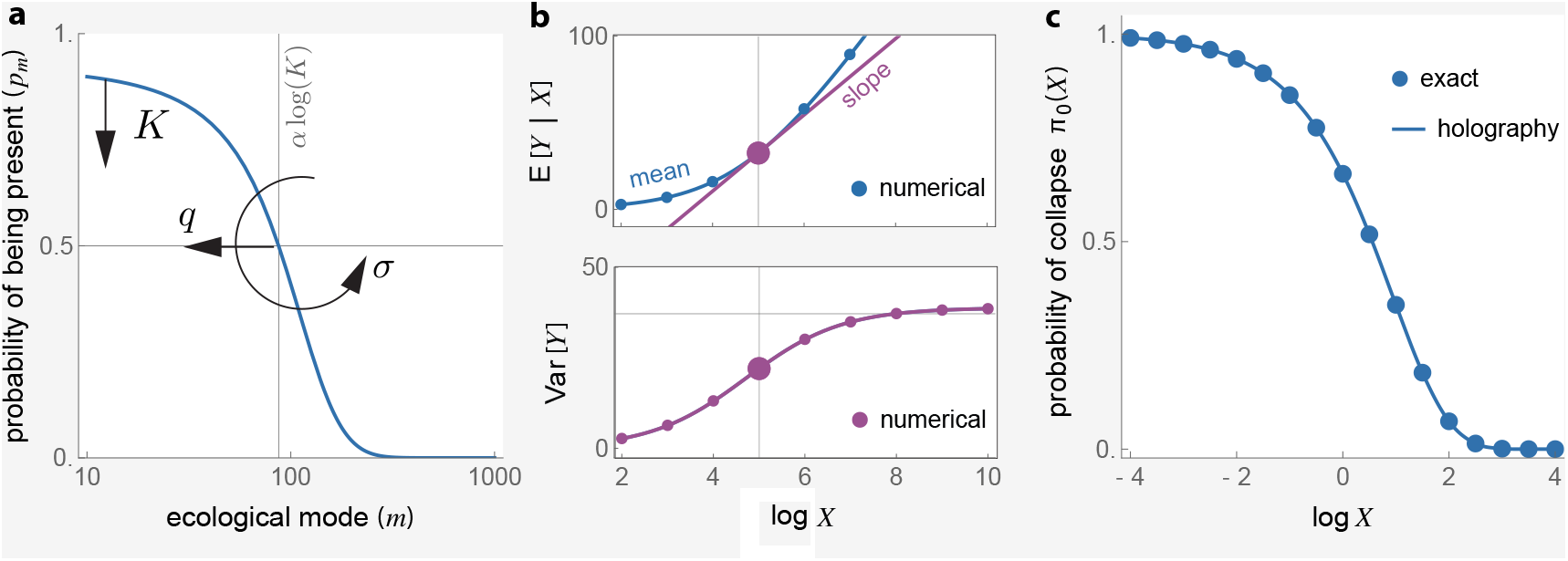

## Methods

### Fisher’s log-series distribution

When the species pool is distributed according to Fisher’s log-series, the number of species with an abundance of *n* individuals is expected to be

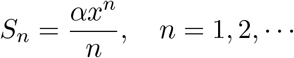

where *α >* 0 is a diversity parameter (called Fishers alpha), controlling the overall richness of the community, and *x*∈(0, 1) is a parameter related to the proportion of rare vs. common species.

### Probability of a new species

We assume individuals of all species in the pool have equal chance of immigrating into the patch. This implies that the probability that an individual of species *i* enters the patch is proportional to the species abundance. We conceptualize this process as choosing at random an individual from outside the patch and letting immigrate. Suppose that after *k* of such random samples we have observed individuals of *Y*_*k*_ different species at the patch. Note this is equivalent to choosing a sample of *k* individuals an observing *Y*_*k*_ different species. Let *p*_new_ = ℙ [*Y*_*k*+1_ *> Y*_*k*_] denote the probability that the next individual belongs to a new species. Under Fisher’s log-series, a classic result that can be traced to Ewens [26] is that

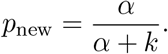

Since *k* is not known to us, our goal now is to provide an estimate of *p*_new_ based on *Y*_*k*_ only. Since *Y*_*i*+1_ − *Y*_*i*_ = **1**{new at *i* + 1}, we obtain

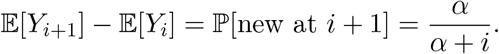

Summing from *i* = 0 to *i* = *k* − 1 yields

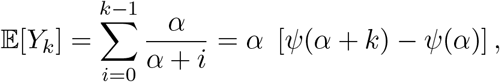

where *ψ* is the digamma function. Treating *Y*_*k*_ as the observed value of this mean yields *Y*_*k*_ = *α* [*ψ*(*α* + *k*) − *ψ*(*α*)]. Solving for *k* we obtain the estimate

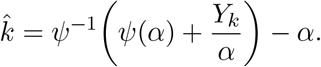

Plugging this estimator in *p*_new_ we obtain

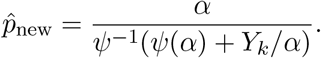

To simplify this formula, we use the asymptotic approximation of the digamma function *ψ*(*z*) ≈log(*z*). Using this approximation we obtain

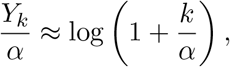

which implies 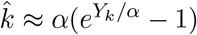. Substituting this in *p*_new_ yields

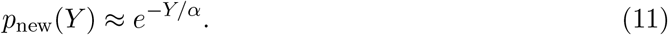

### Species arrival rate

Following previous works, we consider that the immigration rate to a patch of area *X* scales allometrically as *ρX*^*γ*^ individuals per unit of time [18, 19]. Here *ρ >* 0 is the immigration rate to a patch of unit size, and *γ >* 0 is the allometric exponent. This scaling reflects the fact that immigrant influx is proportional to the effective size of the target region, which grows as a power of area depending on whether arrivals are bulklimited (*γ* = 1) or boundarylimited (*γ* = 1*/*2). In general, *γ* thus captures the dimensionality of the process, ranging from areabased deposition to edge or perimetercontrolled immigration.

The rate *λ* of arrival of new species to the patch is this immigration rate multiplied by the probability *p*_new_ that a individual that arrives is from a new species. Using Eq. (11) yields

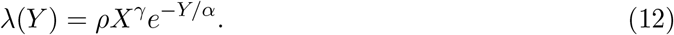

### Species extinction rate

Let *µ >* 0 denote the per-capita mortality (turnover) rate. The extinction rate is the mortality rate times the number of singleton species. From the log-series distribution, the number of singletons is *S*_1_ = *αx*. If the number of species is *Y*, the log-series implies

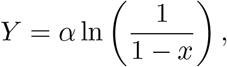

from which we deduce *x* = 1 − *e*^−*Y/α*^. Therefore, the extinction rate is

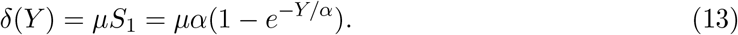

### Deriving the stationary distribution

We model species richness *Y*_*t*_ ∈ {0, 1, 2, …} as a continuous-time birth-death chain with nearest-neighbor transitions. The immigration (richness-increasing) rate from state *y* is *λ*(*y*) and the extinction (richness-decreasing) rate isδ(*y*) as defined in Eqs. (12) and (13), respectively.

Let *π*(*y*) denote a stationary law. Enforcing detailed balance between adjacent states *π*(*y*) *λ*(*y*) = *π*(*y* + 1)δ(*y* + 1) yields

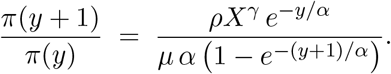

Introduce the emergent parameters

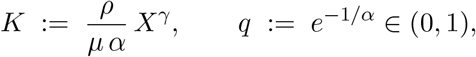

so that *e*^−*y/α*^ = *q* ^*y*^ and 1 − *e*^−(*y*+1)*/α*^ = 1 − *q* ^*y*+1^. Then

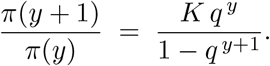

Iterating from *y* = 0 gives

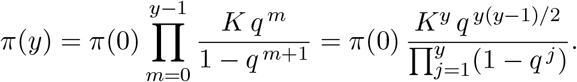

Recognizing the *q*-Pochhammer symbol 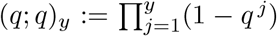 (with (*q*; *q*)_0_ := 1), we obtain the stationary law up to normalization:

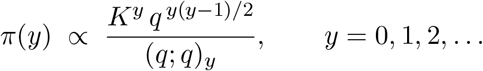

Define the partition function

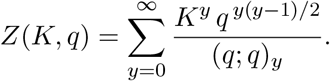

Since the ratio of successive terms is 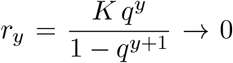 for 0 *< q <* 1, the series converges. By Eulers *q*-binomial identity,

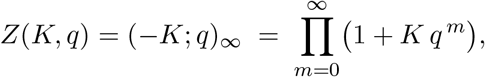

so the normalized stationary distribution is

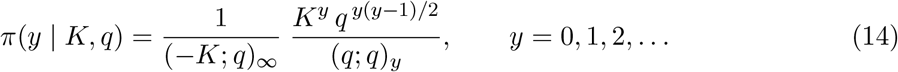

### Ergodicity and entry rates

For any bounded function *f*, the ergodic theorem for continuous-time markov chains implies

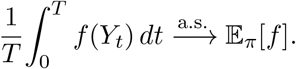

Taking *f* = **1**_*A*_ for any set *A* ⊂ℕ (e.g., *A* = {0} for complete collapse or *A* = {0, …, *y*\*} for very low richness) gives

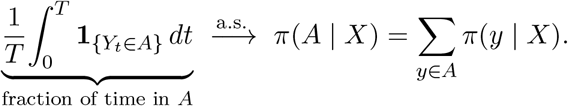

Thus, the stationary tail mass *π*(*A* |*X*) is exactly the point-in-time (ensemble) probability that a patch is in a catastrophic state *A* at equilibrium. In other words, tiny tail probabilities are the frequencies of catastrophic low-richness states you would observe if you sampled the system at a random time once it has equilibrated. In this sense, the stationary law lets us talk rigorously about rare catastrophes.

For a threshold (“catastrophe”) set *A*_*y*_* := 0, 1, …, *y*\*, entries into *A*_*y*_* occur only at the boundary transition *y*\* +1 *y*\*, while exits occur only at *y*\*→ *y*\* →+1. Stationarity imposes equality of boundary flows, hence the *entry intensity* (long-run rate of entries) is

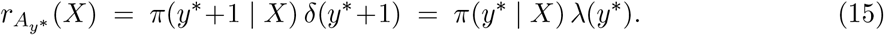

For complete collapse *A*_0_ = {0},

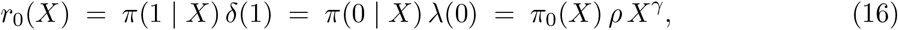

since *λ*(0) = *ρ X*^*γ*^. Writing *S*_*A*_ for the sojourn time in *A* during a single visit, Kac’s renewal identity yields

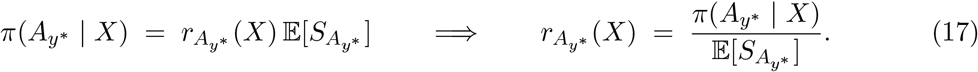

In particular, 𝔼 [*S*_{0}_] = 1*/λ*(0), recovering *r*_0_(*X*) = *π*_0_(*X*) *λ*(0).

### The Fermi-Dirac structure of the stationary distribution

Using Eq. (14), the probability generating function (PGF) of *Y* is

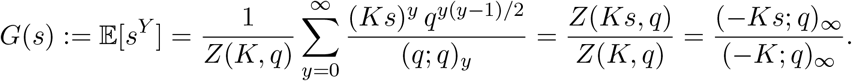

Using the product form of *Z* we obtain

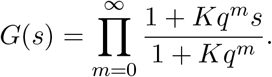

Since 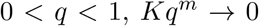 geometrically as *m* → ∞, so the product converges uniformly on |*s*| ≤ 1, and *G* is a valid PGF.

Now define as 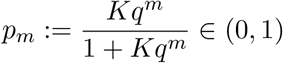. Since 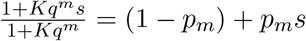, we can rewrite the PGF as

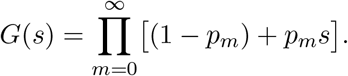

This is the PGF of a sum of independent Bernoulli random variables. This implies that

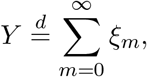

where *ξ*_*m*_ ~ Bernoulli(*p*_*m*_) are independent. Because 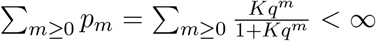, the sum is almost surely finite and all moments exist.

Finally, to derive the Logistic (Fermi-Dirac) occupancies, write *q* = *e*^−1*/α*^ and *K* = *e*^*µ/α*^ with

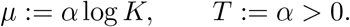

With this notation, 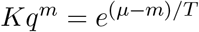 and

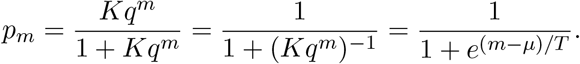

Thus each “ecological mode” *m*∈{ 0, 1, 2, …} is occupied with a logistic probability *p*_*m*_, and richness is the number of occupied modes:

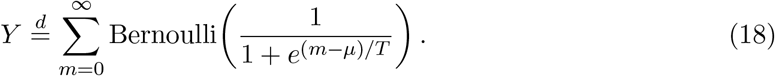

This representation is mathematically identical to the Fermi–Dirac statistics of non–interacting fermions [20, Ch. 3], with *µ* = *α* log *K* playing a biodiversity “chemical potential” and *T* = *α* an effective diversity temperature. This proves the equivalence between the Fermi-Dirac Bernoulli sum and the stationary distribution.

### Scaling laws (Euler-Maclaurin approximations)

Replacing sums by integrals yields:

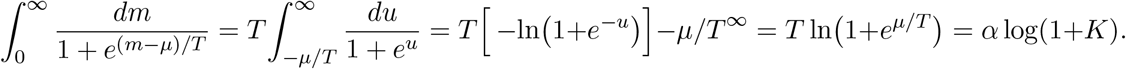

Using *dp/du* = −*p*(1 − *p*) with *u* = (*m* − *µ*)*/T* gives

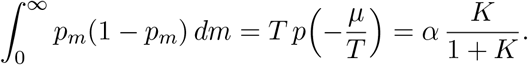

Therefore,

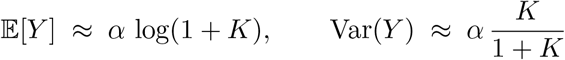

### Corrections in scaling laws due to heterogeneity between patches

To incorporate spatial heterogeneity across plots, we assume that the immigrationextinction ratio *κ* varies lognormally, log *κ*_*i*_ = log *κ*+*σZ* with *Z*~ 𝒩 (0, 1) and *σ*^2^ ≪1. For each patch, the pure model gives mean richness *m*_0_ (*X*; *κ* _*i*_) = *α* log(1 + *κ_i_X*^*γ*^) and variance 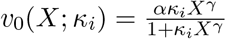 Writing *K* = *κX*^*γ*^ and *v* = *αK/*(1 + *K*), a Taylor expansion around log *κ* yields corrections of order *σ*^2^. The first derivative of *v*_0_ with respect to log *κ*_*i*_ is *v*^′^(*u*) = *αK/*(1 + *K*)^2^ and the second derivative *v*^′′^(*u*) = *αK*(1 − *K*)*/*(1+*K*)^3^. Applying the delta method to the total variance identity 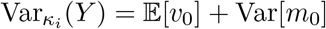 produces an explicit small-*σ* expansion.

Averaging over the random *κ*_*i*_ gives

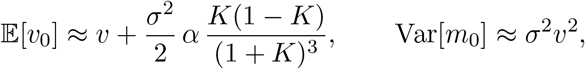

so that

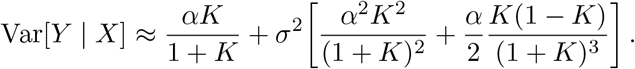

The first term is the Fermi-Dirac variance derived for the homogeneous system, while the bracketed term quantifies the variance inflation due to heterogeneity in *κ*. The correction vanishes both at very small (*K* ≪1) and very large (*K* ≫ 1) areas, where all patches are nearly empty or saturated, and reaches its maximum near *K* ≈1, the ecological Fermi level where richness responds most sensitively to area.

For presentation in the main text, this expression can be simplified to the intuitive rule-of-thumb

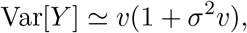

which captures the dominant variance inflation term while preserving the correct asymptotic limits. In this approximation, spatial heterogeneity acts as a multiplicative risk factor that scales with the pure variance *v* = *αK/*(1 + *K*), providing a direct quantitative link between ecological variability (*σ*^2^) and biodiversity risk.

### Risk “free energy” and cumulant identities

The Dirac-Fermi product measure of Eq. (18) defines a grand partition function

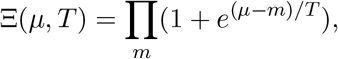

whose logarithm

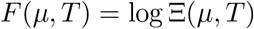

plays the role of a “free energy” in physics. Note that the probability that all modes are empty (i.e., probability of full biodiversity collapse) is

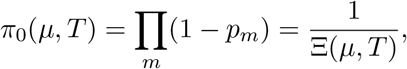

so that

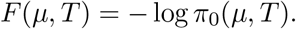

Therefore, *F* can be interpreted as a “risk free energy” and *π*_0_ as its exponential Boltzmann weight.

Furthermore, because the Dirac-Fermi model is an exponential family, derivatives of *F* generate the cumulants of richness:

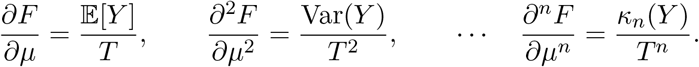

### Scaling with area and the holographic relation

Write the biodiversity potential as *K* = *κX*^*γ*^ with *κ* = *ρ/αµ*. Then *µ* = *α* log *K* = *α*[log *κ* +*γ* log *X*]. Differentiating *F* with respect to log *X* and using *T* = *α* gives

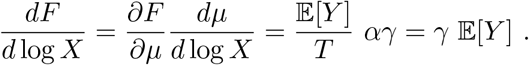

Integration from an infinitesimal area (*X* → 0 ⇒ *F* = 0) yields the identity

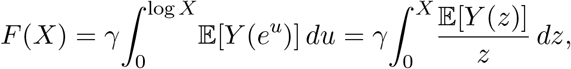

and hence

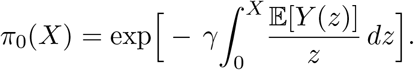

The collapse probability is therefore fully determined by the species–area relation.

### Probability Generating Function (PGF) identity in terms of the mean curve

Let *G*_*X*_(*s*) := 𝔼 [*s*^*Y* (*X*)^] be the PGF of *Y* (*X*). We prove that for any real *s* ∈ (0, 1),

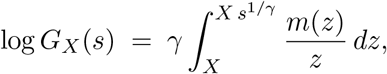

where *s*^1*/γ*^ is taken on the principal branch.

**Lemma**. For any *p* ∈ (0, 1) and *s >* 0,

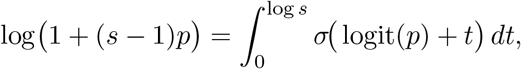

with *σ*(*u*) = 1*/*(1 + *e*^−*u*^) and logit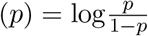. *Proof*. Differentiate both sides with respect to *t* = log *s* and match the value at *s* = 1.

Apply the lemma with *p* = *p*_*m*_(µ(*X*)) and sum over *m*. Since 0 ≤ *σ* ≤ 1 and Σ_*m*_ *p*_*m*_(·) = *m*(·) *<* ∞, dominated convergence justifies interchanging sum and integral:

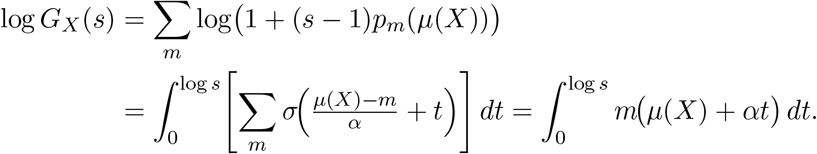

With *µ*(*X*) = *α* log *κ* + *αγ* log *X*, set *u* = *µ*(*X*) + *αt* (*du* = *α dt*), and then *u* = *α* log *κ* + *αγ* log *z* (*du* = *αγ dz/z*). This yields

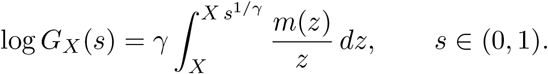

Because both sides are analytic in *s* on |*s*| *<* 1 (principal branch) and agree on (0, 1), the identity theorem extends the equality to all |*s*| *<* 1.

### Biodiversity holography

Let *a*_*k*_(*X*) := Pr{*Y* (*X*) = *k*}. For any 0 *< r <* 1, Cauchys formula gives

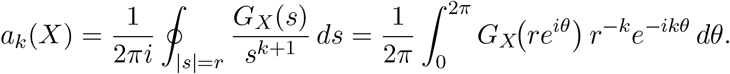

Taking the Abel limit *r* ↑ 1 (standard for power series with nonnegative coefficients) yields

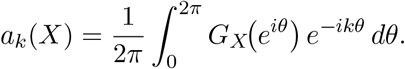

Summing *k* = 0, …, *y* and using 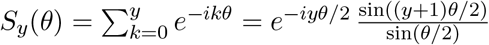.

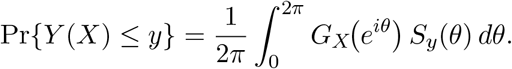

Substituting the PGF identity (by analytic continuation from the real-*s* case to |*s*| *<* 1) with *s* = *e*^*iθ*^ gives

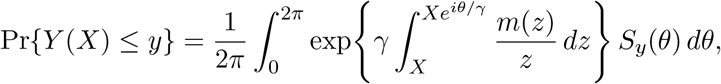

as claimed. In particular, Pr { *Y* (*X*) = 0} = *G*_*X*_(0), and taking the Abel limit *s* ↓ 0 along the positive real axis in the PGF identity yields

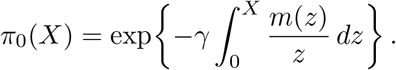

### Convexity and fragmentation penalty

A second derivative yields

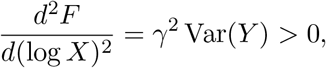

showing that *F* is strictly convex in log *X*. For a landscape divided into patches of area *X*_*i*_ with *θ*_*i*_ = log *X*_*i*_ and mean 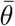, Jensen’s inequality gives

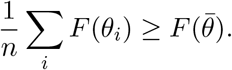

Expanding around 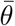 provides the quadratic fragmentation penalty

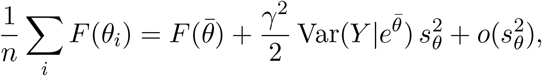

where 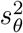 is the variance of log *X*_*i*_. This quantifies the increase in collapse free energy—and hence risk—due to patch-size heterogeneity.

## Data and inference

For each ForestGEO plot, we estimated the first two moments of local richness directly from the census by spatial resampling. Figure 2 uses census 6 for Luquillo, census 1 for CSBI, and census 2 for BCI; other census years yield qualitatively similar results. For our analysis, we considered only living stems with DBH ≥10 cm. For each area *X*, we randomly sampled non-overlapping rectangles of area *X* within the plot and computed richness *Y* in each rectangle. The finite extent of each plot imposes a trade-off between scale and replication: the number of rectangles ranges from ~ 14,000 at small *X* to as few as 3 at the largest *X*. We then estimated the mean richness *m*(*X*) = 𝔼 [*Y* | *X*] and variance *v*(*X*) = Var[*Y* | *X*] from these samples, and used a bootstrap (1,500 resamples) to construct 95% confidence intervals for both moments. As expected from the replication structure, uncertainty is largest for *m*(*X*) at small *X* (high intrinsic variability across many small patches) and for *v*(*X*) at large *X* (few independent rectangles).

We inferred model parameters by exploiting the predicted asymptotic scaling of the first two moments (Box 1, point 1). We first identified a large-area regime and fit a linear model in (log *X, m*) using Mathematicas `LinearModelFit`. The fitted slope estimates the product *αγ*. We then identified a small-area regime and fit a linear model in (log *X*, log *m*). The slope estimates *γ*, and the intercept at *X* = 1 estimates log(*ακ*) (equivalently, *ακ* on the original scale). These three constraints determine *α, γ*, and *κ*. We identified the small-area (resp. large-area regimes as those for which the inferred parameters changed less than 10% when the area decreased (resp. increased). Finally, holding *α, γ*, and *κ* fixed, we inferred the mixing parameter *σ* from the analytical expression for *v*(*X*) using Mathematicas `NonlinearModelFit`. Parameter estimates are summarized in Table 1.

## Acknowledgments

This research was supported by grants BSR-8811902, DEB 9411973, DEB 0080538, DEB 0218039, DEB 0620910, DEB 0963447 AND DEB-129764 from NSF to the Department of Environmental Science, University of Puerto Rico, and to the International Institute of Tropical Forestry, USDA Forest Service, as part of the Luquillo Long-Term Ecological Research Program. The U.S. Forest Service (Dept. of Agriculture) and the University of Puerto Rico gave additional support. The LFDP is part of the Smithsonian Institution Forest Global Earth Observatory, a worldwide network of large, long-term forest dynamics plots. Funding for the Smithsonian Conservation Biology Institute (SCBI) Large Forest Dynamics Plot (LFDP) was provided by the Smithsonian Institution, the National Zoological Park, and the HSBC Climate Partnership. The SCBI LFDP is part of the Smithsonian Institution Forest Global Earth Observatory, a worldwide network of large, long-term forest dynamics plots.

## Data accessibility

Data analyzed in this work is available from the ForestGEO database.

## Generative AI and AI-assisted technologies in the writing process

During the preparation of this work, the author used ChatGPT in order to improve language and readability. After using this tool/service, the author reviewed and edited the content as needed and takes full responsibility for the content of the publication.

---

**Supplementary Figure 1:**
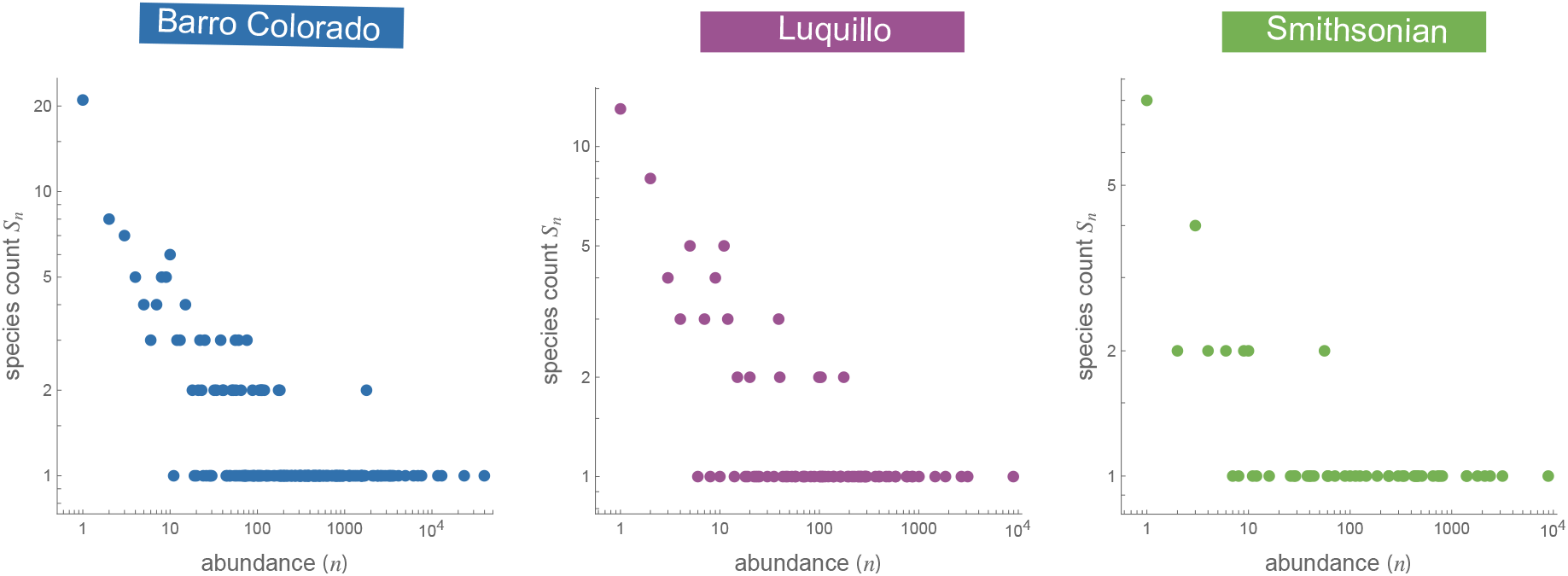
Empirical speciesabundance counts-of-counts across ForestGEO plots. For each plot (Barro Colorado Island, Luquillo, and Smithsonian Conservation Biology Institute), we used the full census of living stems with DBH ≥ 10 cm and summarized the speciesabundance distribution as *S*_*n*_, the number of species represented by exactly *n* individuals. Points show the observed pairs (*n, S*_*n*_) on logarithmic abundance axis. The strong excess of rare species and the long right tail are characteristic of Fisher-type log-series structure and motivate the regional log-series approximation used in the theory.

